# *In vivo* fate of free and encapsulated iron oxide nanoparticles after injection of labelled stem cells

**DOI:** 10.1101/366518

**Authors:** Sumaira Ashraf, Arthur Taylor, Jack Sharkey, Michael Barrow, Patricia Murray, Bettina Wilm, Harish Poptani, Matthew J. Rosseinsky, Dave Adams, Raphaël Lévy

## Abstract

Nanoparticle contrast agents are useful tools to label stem cells and monitor the *in vivo* bio-distribution of labeled cells in pre-clinical models of disease. In this context, understanding the *in vivo* fate of the particles after injection of labelled cells is important for their eventual clinical use as well as for the interpretation of imaging results. We examined how the formulation of superparamagnetic iron oxide nanoparticles (SPIONs) impacts the labelling efficiency, magnetic characteristics and fate of the particles by comparing individual SPIONs with polyelectrolyte multilayer capsules containing SPIONs. At low labelling concentration, encapsulated SPIONs served as an efficient labelling agent for stem cells. The bio-distribution after intra-cardiac injection of labelled cells was monitored longitudinally by MRI and as an endpoint by inductively coupled plasma-optical emission spectrometry. The results suggest that, after being released from labelled cells after cell death, both formulations of particles are initially stored in liver and spleen and are not completely cleared from these organs 2 weeks post-injection.

## Introduction

Stem cell regenerative medicine therapies have been proposed for the treatment of a range of debilitating conditions.^1-4^ Tracking of stem cells in pre-clinical models is a prerequisite to determine their safety and efficacy.^5-7^ For this purpose, in addition to, and/or in combination with genetic reporters, nanoparticles are successfully used as contrast agents for cell labelling and tracking with various imaging modalities.^8-12^

Cellular biodistribution, viability and proliferation can be monitored by bioluminescence imaging (BLI) the cells transduced with the genetic reporter firefly luciferase (Luc).^12-14^ Cell death results in a decrease in bioluminescence signal, whilst cell division and tumor formation leads to an amplification of the signal. This is a highly sensitive and robust method of monitoring the biodistribution of cells though it has a low spatial resolution (intra and inter organ monitoring of cells distribution is not possible).^15^

Magnetic resonance imaging (MRI) offers much higher spatial resolution (50 µm) than BLI allowing *in vivo* cell tracking combined with detailed anatomical information at the level of individual organs.^16^ Appropriate MRI contrast agents^17, 18^ for cell tracking include superparamagnetic iron oxide nanoparticles (SPIONs) which generate a negative contrast.^11, 19-24^ The uptake efficiency of SPIONs is strongly influenced by their functionalization.^21, 25^ Recently, positively-charged DEAE (diethylaminoethyl)-dextran coated SPIONs have been synthesized for enhanced cellular uptake and MRI contrast.^20, 22^ Upon cell uptake, the clustering and confinement of these particles in endosomal and lysosomal compartments affects their abilities to alter the relaxation rate and hence relaxivity of the surrounding water molecules.^26, 27^ The critical parameter for imaging is the contrast obtained after cell labelling rather than the solution relaxivity of the particles.^20, 22^

One potential approach to enhance SPIONs uptake by cells, is to assemble them inside polymeric capsules resulting in the packing of large amount of individual nanoparticles within a confined volume.^28-30^ Polyelectrolyte multilayered (PEM) and multifunctional capsules are fabricated by layer-by-layer (LbL) assembly of oppositely charged polymeric layers around a template.^31^ Their design can be tailored by multiple strategies and particle loading can be enhanced by including more layers of particles during the capsule’s assembly.^32^ The choice of the final polymer layer determines their cellular interaction, uptake efficiency and hence internalization. It has been observed that a final layer of positively charged polymers results in increased uptake of capsules.^33^ For long term cellular imaging the walls of the capsules can be made of non-biodegradable materials.^34^

We and others have shown that most cells die after injection.^11, 15, 35^ Therefore, understanding the *in vivo* fate of cell-labelling nanoparticles, especially following the death of the labelled cells is important for the interpretation of imaging studies and to assess the risk of toxicity. In the current study, we focus on the effect of formulation on the fate of SPIONs after the *in vivo* injection of labelled cells. More specifically, we labelled mouse bone marrow derived mesenchymal stem cells (mMSCs) with free and PEM-encapsulated SPIONs. First the solution relaxivities, magnetic properties, toxicity, and cell labelling efficiency of both formulations of particles are compared. Then, after intra-cardially (IC) injecting the labelled cells in the left ventricle of mice, the animals were longitudinally imaged by BLI and MRI until 14 days post injection. The cellular bio-distribution and viability was monitored by BLI, whilst the MRI allowed visualization of the *in vivo* fate of particles after cell death. At the end point of experiments (2 weeks post-injection), the amount of elemental iron inside the mice organs was measured to analyze the accumulation and elimination of both formulations of SPIONs. Our studies demonstrate a similar accumulation and elimination pattern of particles injected via labelled cells to what has been reported for the particles directly injected in the blood stream.^36-40^

## Results and Discussion

### Preparation and characterization of free and encapsulated SPIONs

To evaluate the effect of incorporation of the SPIONs into capsules, magnetization curves were acquired for capsules and free particles; no difference was observed between the magnetic properties of the free and encapsulated SPIONs (Fig. 1). Whilst the properties of individual particles were not affected by encapsulation, the large number of particles in a single object resulted in higher magnetic forces that enabled magnetic separation of particles during purification from excess reactants. Thus, during LbL assembly of particles in polymeric capsules, as the number of particles per capsule increased, their magnetic separation by a bar magnet became possible. After deposition of 3 layers of SPIONs, for further layer deposition, the excess reactants were therefore removed by magnetic separation instead of centrifugation. The transmission electron micrographs (Fig. 1 a & b) show that particles did not fuse inside the capsules; they retained their identity as individual particles (as indicated by the magnetization curve; Fig. 1 c). The crystalline structure and particle size (~ 8 nm) was confirmed by powder XRD (pXRD) before encapsulation (Fig. SI.1 a). Their hydrodynamic diameter in water was measured by dynamic light scattering (DLS, ~85 nm, Fig. SI.1 b). The hydrodynamic diameter of capsules cannot be measured accurately by DLS because of their dimension (µm size) and sedimentation behavior.

**Figure 1:**
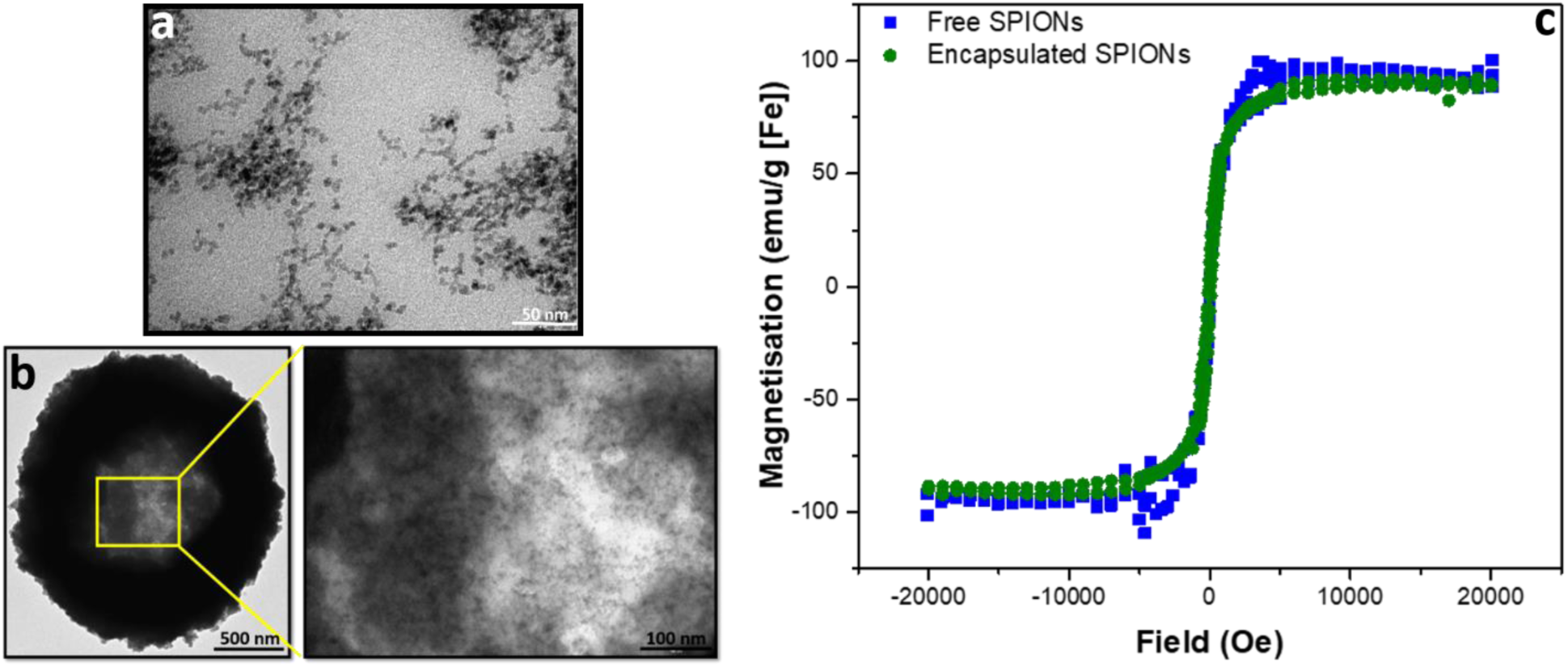
Characterization of free and encapsulated SPIONs by transmission electron microscopy (TEM) and superconducting quantum interference device (SQUID). TEM of free and encapsulated SPIONs is presented in (a) and (b), respectively. SQUID magnetisation curves are shown in (c).

### Effect of encapsulation of particles on the solution relaxivity

To determine the effect of encapsulation on the solution relaxivity, the water relaxation rates of particles at different concentrations (based on Fe content) were measured by MRI and plotted as function of concentration (Fig. 2). Encapsulation of SPIONs resulted in a drop in solution relaxivity due to hindered access to the solution water protons and is similar to the previous findings of the entrapment of free particles inside the lysosomes.^20, 22, 32^

**Figure 2:**
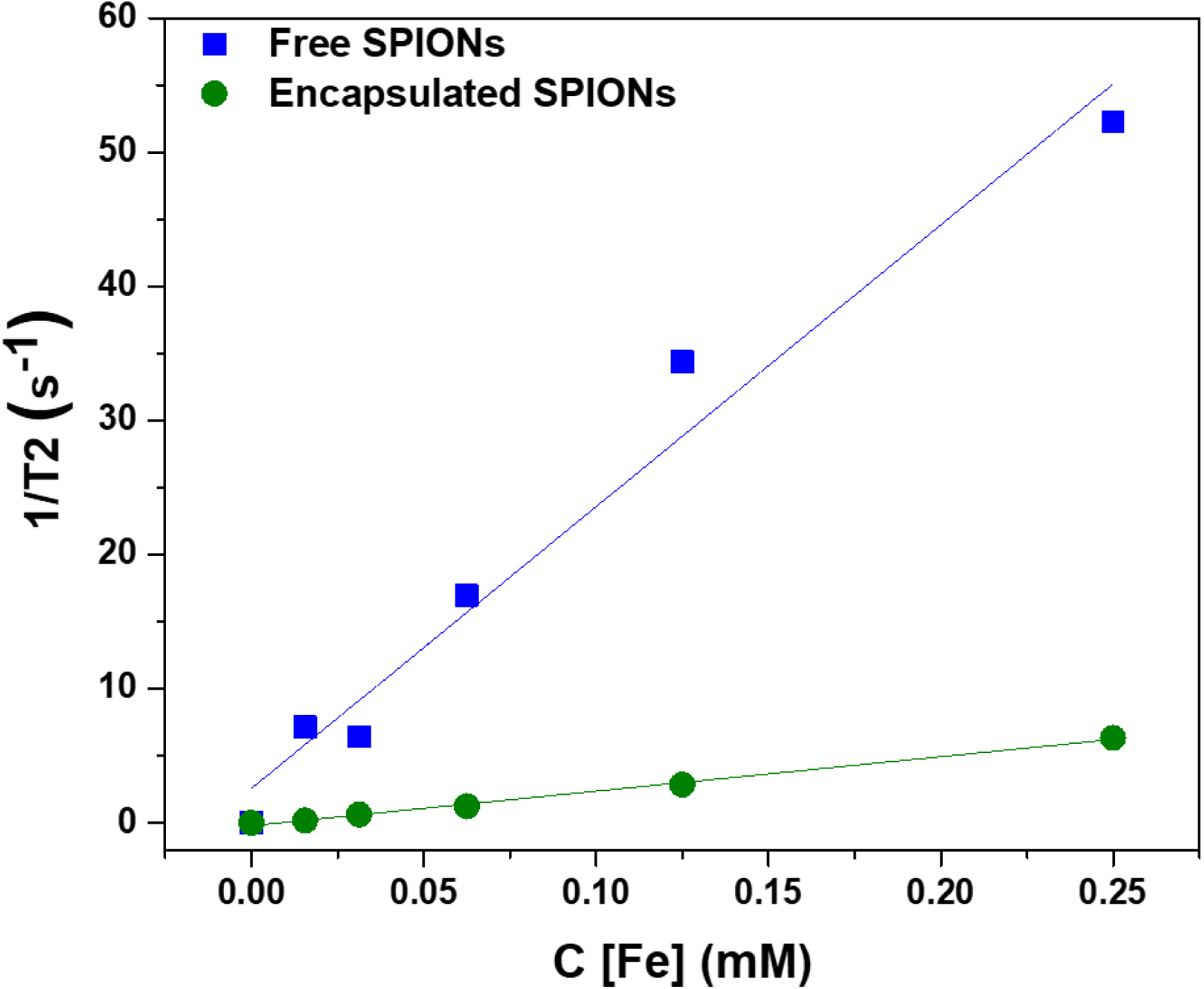
Effect of encapsulation of SPIONs on the solution relaxivity. The water relaxation rates are plotted against the concentrations of particles (in terms of Fe content).

### Cell labelling efficiency and toxicity of SPIONs

mMSCs were chosen for this study because they are similar to human bone marrow derived mMSCs which are being used in clinical trials,^41, 42^ and it has already been shown that these cells uptake free SPIONs and other nanoparticle probes with good efficacy.^12, 22^ To evaluate the cell labelling efficiency and toxicity of free and encapsulated SPIONs, cells were labelled with suspensions of particles (encapsulated or free) having comparable Fe content. The cells were imaged by MRI to quantify the solution relaxation times (ms; Fig. 3 a & b) and the Fe content in labelled cells was determined with a ferrozine assay (Fig 3, c). This revealed a two-fold higher Fe uptake by the cells when labelled with encapsulated SPIONs as compared to the free particles.

**Figure 3:**
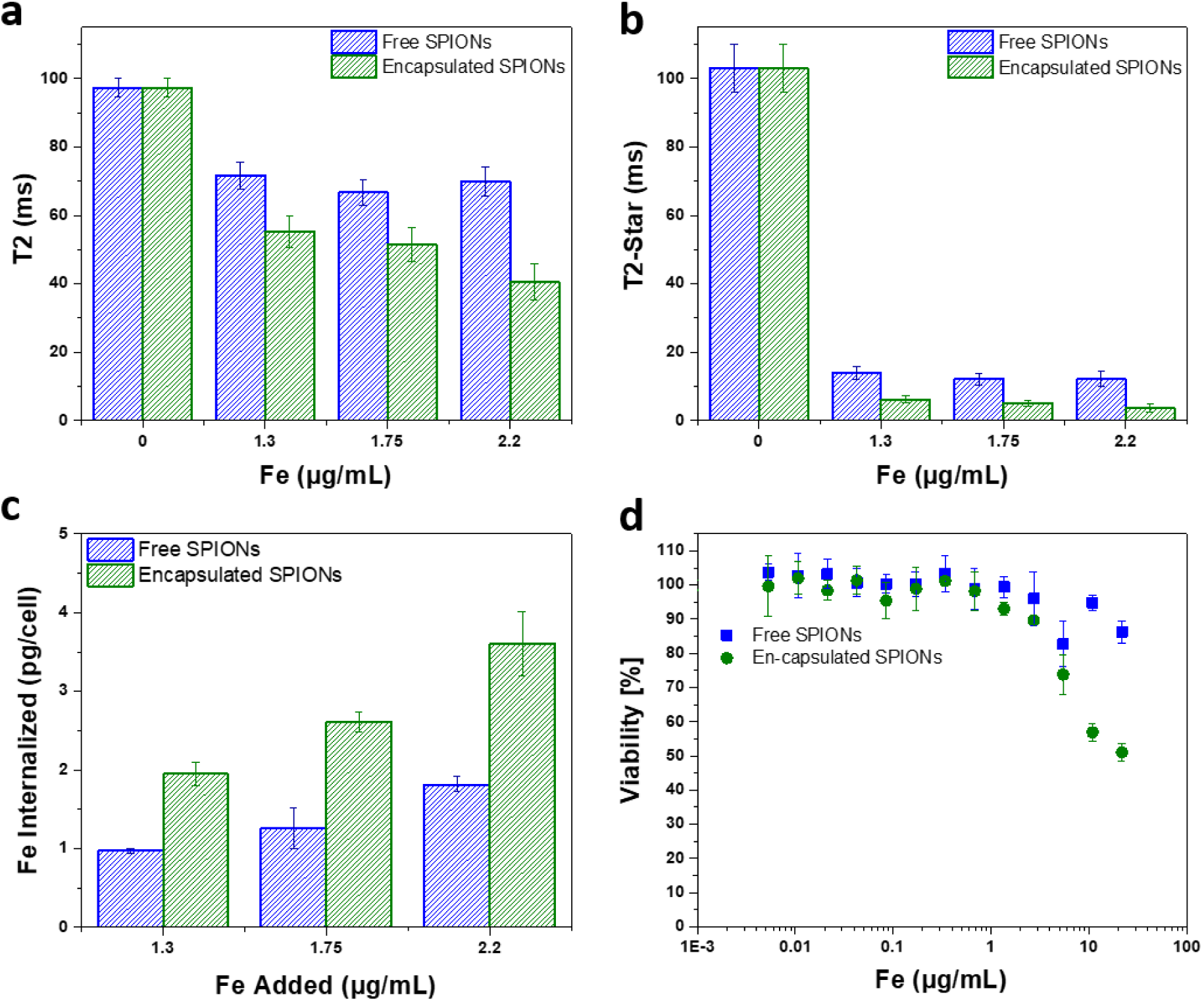
*Cell labelling and toxicity of SPIONs. The solution relaxation times (T2 and T2-star) of the particles are plotted against the concentrations used for cell labelling (a & b)*. The labelling concentrations of Fe (Fe added to label the cells) are presented along abcisa. *Internalization of free and encapsulated SPIONs versus the added concentration of particles used for cell labelling (determined from ferrozine assay) are shown (c). The Fe concentrations of 1.3, 1.75, and 2.2 µg/mL corresponds to the doses of 15, 20, and 25 capsules added per cell. Cell viability against the labelling concentrations of Fe is presented (d). The error bars are the standard deviations of three replicates*.

The difference in Fe uptake is unlikely to be due to electrostatic interactions,^25, 33^ as both formulations were highly positively charged (Fig. SI.1 c & d). However, zeta potential measurements for large sedimenting objects (encapsulated SPIONs) is not reliable as sedimentation during the measurement period affects the accuracy of the data. Due to the large number of SPIONs per capsule, the uptake of a few capsules per cell resulted in increased Fe uptake per cell (Fig 3, c). Depending upon the cell line/type and size/dimensions of capsules, the number of micro capsules phagocytosed by cells cannot exceed a limit of 2 - 15 capsules/cell.^33^ So for low labelling concentrations a higher Fe uptake is achieved by using encapsulated particles. Higher degrees of labelling have been reported with high labelling concentrations of free particles.^19^ In addition to efficient labelling at low Fe concentrations, capsules can be used to combine nanoparticles in a single entity and hence can facilitate multiplexed imaging and measurements.^43^

The toxic effect of free and encapsulated particles is comparable except at high doses of 125 and 250 capsules added per cell (Fig. 3 d). This may be due to the sedimentation of the excessive encapsulated particles (µm dimension; all particles cannot be endocytosed) resulting in the formation of a layer entirely covering the cells’ surface leading to very high local concentration. In the following sections, the cells were labelled with particles below their toxicity level. For *in vivo* experiments, 15 capsules per cell (1.3 µg [Fe]/mL) and ~2.6 times higher concentration of free particles (3.3 µg [Fe]/mL) were added to the cells to obtain a similar amount of iron per cell (~4 and ~5 pg Fe/cell, respectively).

### Bioluminescence of mMSCs

To verify the mMSCs Luc^+^ bioluminescence signal, D-luciferin was added to the cells and the bioluminescence signal was detected after 15 min of incubation using an IVIS spectrum imaging system. It was noticed that the bioluminescence signal from ~200 cells was detectable (Fig. SI.2), indicating that the intensity of the bioluminescence signal was sufficient for *in vivo* detection.

### Cell phenotype after labelling and re-seeding/re-spreading

To observe cell phenotype, the trypsinized cells suspended in ice cold PBS for 5 – 6 hrs were plated (re-seeded). Three days post-seeding the morphology of the labelled cells observed by light microscopy was not distinguishable from unlabeled cells, irrespective of whether they were labelled with free or encapsulated SPIONs (Fig. SI.3). Capsules can be seen inside the labelled cells (Fig. SI.3).

### Animal injection and in vivo monitoring of the bio-distribution of labelled cells

When cells are injected into mice, a significant proportion of them die within a short time-frame.^5, 42, 44^ Due to the impact on the interpretation of imaging results and particles’ associated toxicity, it is important to see the fate of particles once labelled cells die. Hence, soon after injecting the labelled mMSCs into mice, their *in vivo* bio-distribution was monitored in parallel by BLI and MRI for 14 days. As expected, the vast majority of the cells were no longer detectable by day 1. Surprisingly, in some animals, a strong luminescence signal was detected in the cardiac regions throughout the course of imaging (Fig. 4); this was likely due to small numbers of cells from the needle tract engrafting in the cardiac muscles following IC administration. By day 14, luminescence signals were present in the hind regions of some mice. As we have previously shown, these were likely to be osteosarcomas.^11^ On culling the mice, no tumor was detected in the major body organs (liver, spleen, brain, heart, and lungs). A key objective of this study was to investigate what happened to the free and encapsulated SPIONs after day 1 when BLI showed that the majority of the cells had died. For this purpose, we require consecutive MRI and BLI to be performed on the same animals.

**Figure 4:**
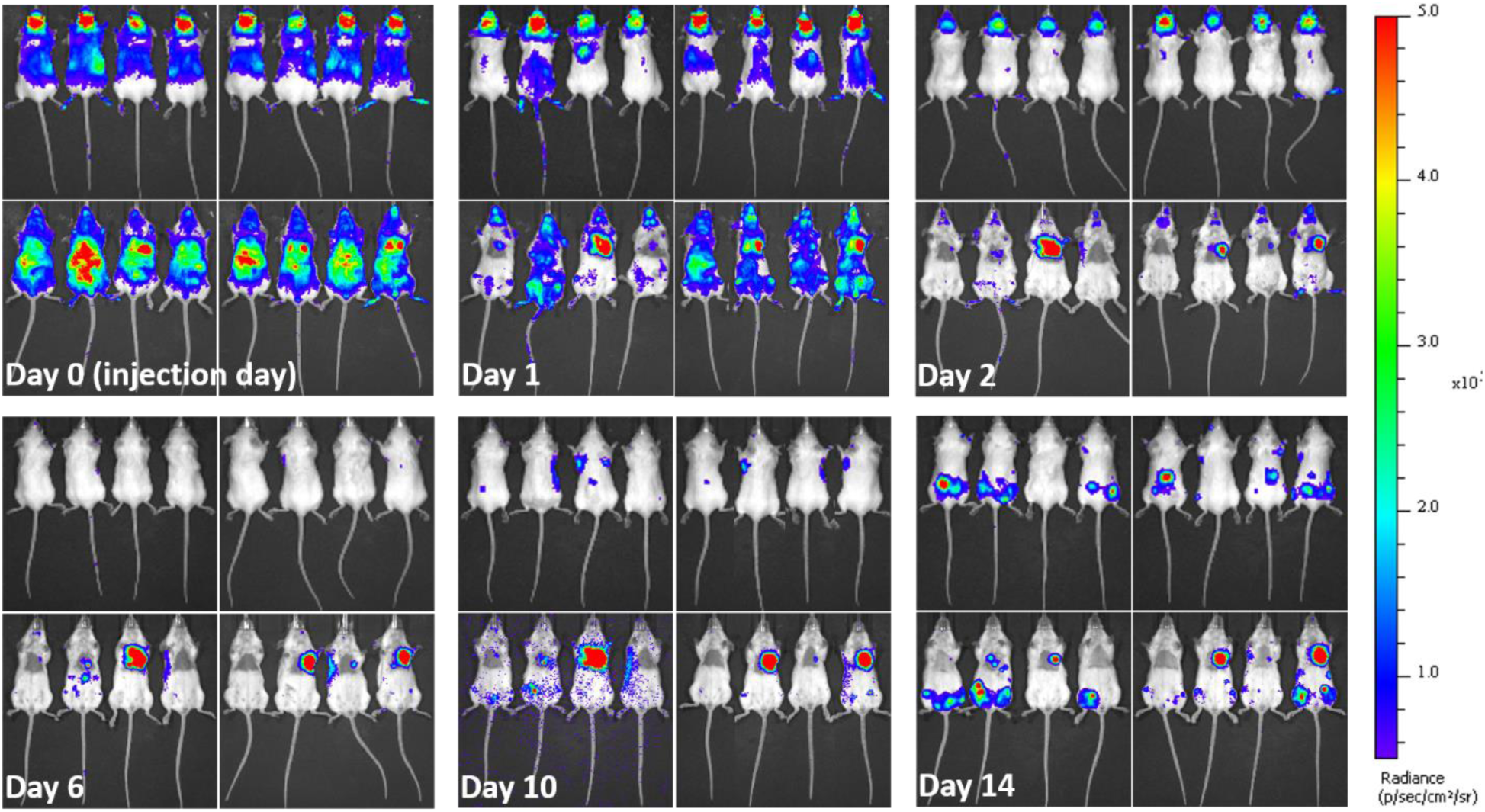
In vivo long term fate of labelled mMSCs determined from bioluminescence imaging. Images taken at different days post injection of labelled cells. Left panels in each image set (each day) represent mice injected with mMSCs labelled with free SPIONs. While, right panels represent mice injected with mMSCs labelled with encapsulated SPIONs. Upper and lower rows show dorsal and ventral aspects, respectively. Images were recorded 15 minutes after intraperitoneal (IP) injection of D-luciferin.

### MRI complementary to BLI

The high resolution of MRI facilitates the visualization of the intra-organ bio-distribution of the contrast agent.^11^ Here, MRI allowed the monitoring of *in vivo* clearing and accumulation pattern of particles (Fig. 5 and SI.4), after confirmation of cell death by BLI. Since the major Fe accumulation and clearing/elimination organs are kidneys, liver, and spleen, we imaged the entire abdominal region of mice by MRI throughout the course of experiments to assess the presence and distribution of the particles. Significant difference in contrast can be seen in the images over the time-course of the experiment, being highest soon after injection and reducing towards baseline over the following days (Fig. SI.4). Full MRI datasets are available on the Zenodo data archive; https://doi.org/10.5281/zenodo.1203991.

**Figure 5:**
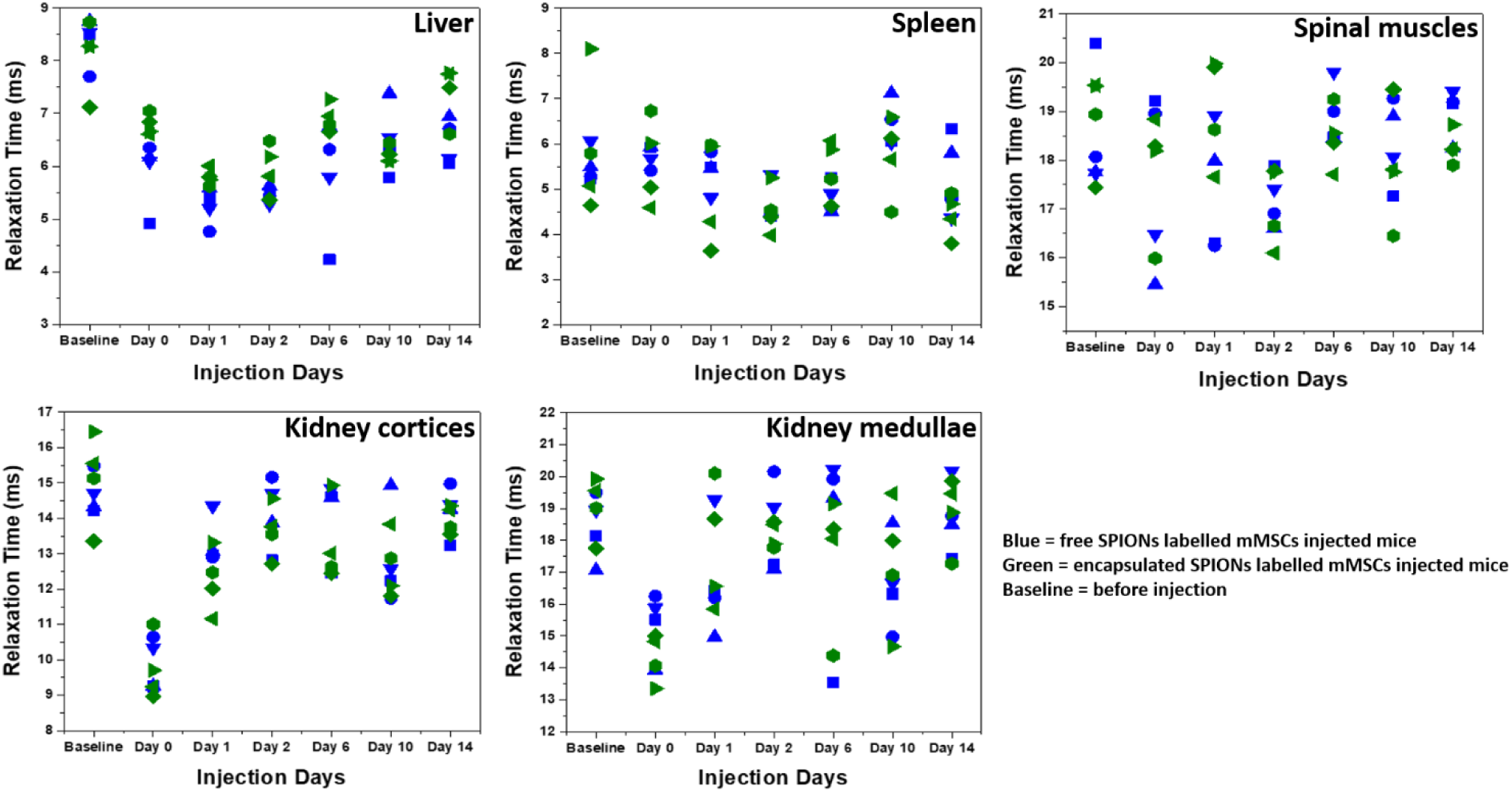
Relaxation times (T2-Star) as function of time. T2* relaxation times of SPIONs in different organs and tissues are plotted at different days. Different symbol shapes represent different animals.

To quantify the Fe accumulation and clearance pattern before and after injection, the relaxation times (ms) in some organs and tissues (liver, spleen, spinal muscles, kidney cortices and medullae) were calculated (Fig. 5) from the MRI scans by drawing regions of interests (ROIs) around these organs and tissues. Soon after injection, the relaxation time decreased in the liver for both free and encapsulated SPIONs and a further decrease can be seen in the following days post-injections. The BLI data indicates that 24 hours post-injection there was considerable cell death. Therefore, the decrease in relaxation time in the liver on day 1 and 2 post-injection is possibly due to SPIONs accumulating in the liver following their release from cells. The presence of SPIONs in the liver up to 48 hours post administration is in accordance to the recent findings of Scarfe *et al*.^11^ On day 6 post-injection, the relaxation time in the liver started to increase but remained below the baseline value during the entire time course of imaging (Fig. 5). This suggests that the SPIONs accumulated in liver were not completely cleared/eliminated by day 14. The tendency of accumulation and hence slow clearance of SPIONs from the liver of mice after cell death is similar to the fate of free particles systemically injected in mice.^36^

The spleen has very short relaxation time baseline values, and no further reduction was observed until the last imaging day in the case of free SPIONs (Fig. 5). By contrast, relaxation times appeared to be lower for encapsulated SPIONs at day 14 post injection. Interestingly, the relaxation times for encapsulated SPIONs had increased in the liver at this time point, suggesting that after leaving the liver, encapsulated particles accumulated inside the spleen. However, detailed biochemical pathways for Fe accumulation in these organs need to be explored. Recently, the long term *in vivo* fate of different hybrids of SPIONs with gold nanoparticles has been established after directly injecting the particles in the blood stream of mice which revealed a predominant accumulation of particles in liver and spleen for longer periods of time.^36^ In similar studies, the accumulation and metabolism of particles in the liver and spleen and loss of magnetic characteristics as a function of their surface coating, inner core and formation of protein corona was evaluated.^37-40^ The electron paramagnetic resonance (EPR) based measurements of iron oxide nanocubes showed their considerable elimination from liver on day 7 post administration, whereas half of the proportion of particles stored in the spleen was still detected on day 7 post injection.^39, 40^ In general, free SPIONs have faster lysosomal degradation and cellular elimination rate as compared to the aggregated particles.^45^

The relaxation time and MRI contrast in spinal muscles did not change considerably throughout the time course of experiments (Figs. 5 and SI.4). The SPIONs detected in the kidney cortices and medullae immediately after administration were mostly cleared by day 1 post injection (Fig. 5). This finding suggests that after cell death, the particles were rapidly cleared from the kidneys, in line with what has been reported for the intravenous and IC injected particles where particle accumulation inside kidneys is not observed.^11^

### Inductively coupled plasma-optical emission spectrometry (ICP-OES) based Fe quantification

To confirm the results of the MRI analysis, at the end point of imaging experiments the animals were culled and *ex-vivo* ICP-OES based Fe quantifications in 6 organs (brain, kidney, liver, lung, spleen & heart) were performed (Fig. SI.5). This analysis did not show a significant difference in the percentage of Fe per dry weight of organs at day 14 post injection from the control group of mice which did not receive any injection. This suggests that the majority of the particles released on cell death were cleared by 2 weeks. Although the sensitivity of such measurements is limited as the presence of high quantities of endogenous Fe and animal-to-animal variability renders their interpretation difficult. More elaborate measurement strategies, e.g. isotope labelling,^37, 46, 47^ could be employed in future studies.

## Materials and Methods

### Materials and Reagents

Diethylaminoethyl (DEAE)-dextran (Mw 40 kDa, #80881), iron (III) chloride hexahydrate (#1.03943), iron (II) chloride tetrahydrate ( 98%, #380024), ammonium hydroxide solution (28 – 30 %, # 221228), sodium azide (#S2002), calcium chloride dihydrate (#223506), sodium carbonate (#S7795), sodium chloride (#S7653), poly(sodium 4-styrenesulfonate) (PSS, M^w^ ~70 kDa, #243051), dextran (average Mw 1500 - 2800 kDa, # D5376), ethylenediaminetetraacetic acid disodium salt dihydrate (EDTA, #E5134), sterile dimethyl sulfoxide (DMSO, #D2650), Sephadex^®^ G-100 beads (#GE17-0060-01), 3-(2-pyridyl)-5,6-di(2-furyl)-1,2,4-triazine-5’,5”-disulfonic acid disodium salt (ferrozine reagent, #82940), 2-hydroxyethyl agarose (low gelling temperature, #A4018 and #A9414), sucrose (#S7903), L-ascorbic acid (#95209), ammonium acetate (#A1542), neocuproine (#N1501), and Corning^®^spin-filters (100 kDa, # CLS431491), were purchased from Sigma Aldrich. Spectra/pore biotech cellulose ester (CE) dialysis membranes (MWCO: 100 kDa, # 11415949) and Millex GP syringe filters with polyethersulfone (PES) membranes (0.22 µm, #10038041) were purchased from Fisher Scientific, UK. Membrane filters (#SLHAO33SS; pore size 0.45 µm, and #780532; 0.22 µm) were from Millex HA Millipore and STARLAB, respectively. Poly(allylamine hydrochloride) (PAH, M^w^ ~120 - 200 kDa, # 43092.09) was purchased from VWR. Gelatin capsules (size 4, # AGG29214) were from Agar Scientific. Double distilled deionized (Millipore Limited, Hertfordshire, UK) water with resistivity of 18.2 MΩ.cm was used in all experiments.

Luciferase transfected bone marrow derived mouse mesenchymal stem cell line (mMSCs; D1 ORL UVA [D1](ATCC^®^, #CRL-12424™) was generously provided by Dr Antonius Plagge and was a gift from Bryan Welm (Addgene plasmids # 21375 and 39196). For cell culture, Dulbecco’s modified Eagle’s medium (DMEM, #D6546), phosphate buffered saline (PBS; #806544), PBS tablets (#P4417), and penicillin streptomycin (P/S, #P4333) were purchased from Sigma-Aldrich. L-glutamine (#25030081), fetal bovine serum (FBS; #16000044), and 0.5% Trypsin-EDTA (#15400054) were purchased from GIBCO (Life Technologies). Paraformaldehyde (PFA, 16%, # 28908) was from Thermo Fisher Scientific. Cells were counted by an automated cell counter (TC20™) from Bio Rad. CellTiter-Glo^®^ reagent (#G7570) was from Promega. Cell culture petri dishes (PS, 100/20 mm, vents, Cellstar^®^ #664970), and falcon tubes (Cellstar^®^; 15 mL; #188271, and 50 mL; #227261) were from Greiner Bio-one. 24 well plates (#3524) were from Costar (Corning). 100 well transparent (#167008) and opaque bottom plates (#CLS3917) were from Thermo Scientific and Sigma Aldrich, respectively. Eppendorfs (1.5 mL; # 0030120086) were from Eppendorf. 100 (#981103) and 200 µL (#I1402-8100) polypropylene tubes (eppendorfs) were purchased from Qiagen and STARLAB, respectively. Centrifuge machines (Sigma 2-6E) and (Heraeus Pico 21, Thermoelectron Corporation) were used for centrifuging the mMSCs. Formvar/carbon-coated 200 mesh copper grids (#F077/100) were purchased from TAAB.

### Characterization Tools

The hydrodynamic diameter and zeta potential measurements were performed in water using a Malvern zetasizer Nano ZS (dynamic light scattering). According to the manufacturer’s instructions, the ATP assay (Cell Titer-Glo luminescent assay, Promega) based luminescence (for determination of viability of cells) and ferrozine reagent assay based colorimetric changes (for quantification of iron content) were measured with Fluostar Omega (BMG Labtech) coupled with a microplate reader. Transmission electron microscope (TEM) micrographs were taken using a Tecnai G3 spirit TEM at 120 keV. 5 µL droplets of the diluted samples were drop-casted onto the formvar/carbon-coated 200 mesh copper grids and air dried before TEM analysis. The morphology of mMSCs was observed by bright field (Leica light microscope DM IL coupled with Leica DFC 420C camera) microscopy. For ICP-OES Agilent 5110 ICP-OES spectrometer (with SVDV detection and equipped with the sample changer) was used. To determine the crystalline state and mean core diameter of free SPIONs powder X-ray diffraction (pXRD) measurements were carried out on a Panalytical X’pert Pro multipurpose diffractometer (Co Kα source, λ = 1.78 Å, patterns measurement > 20 – 120° 2θ for 2 hours, step size = 0.033°, time/step = 295.3 s, scan speed = 0.014 °/s). Particles samples were freeze-dried using Labconco freezone 4.5 (condenser temperature –50 °C, shelf temperature 20 °C) and their magnetization behaviour (magnetic isotherm) was recorded by means of a superconducting quantum interference device (SQUID) magnetometer (MPMS XL-7, quantum design, USA) at 300 K with maximum applied field 2 T under helium atmosphere. For SQUID measurements, freeze-dried samples (a few mg) were embedded in gelatin capsules (size 4) which were suspended in the middle of plastic drinking straws. Organs were freeze-dried to obtain their dry weight before ICP-OES based quantification of iron. For bioluminescence (BLI) measurements a bioluminescence imager (IVIS Spectrum, Perkin Elmer, UK) was used to see the *in vivo* biodistribution of mMSCs. Magnetic resonance imaging (MRI) was performed with Bruker Avance III spectrometer interfaced to a 9.4T magnet system (Bruker Biospec 90/20 USR) using a 40 mm transmit/receive volume coil for SPIONs phantoms, SPIONs-labelled cell phantoms and animal imaging. Ultrasound system (Prospect) with utrasound pulse sequence (UPS) user interface from S-Sharp corporation Taiwan was used during IC injections of labeled mMSCs to mice.

### Animals

6 - 8 week old 11 female BALB/c mice were obtained from Charles River, UK and housed in groups of 4 with *ad libitum* access to standard food and water (ventilated cages, 12 h light/dark cycle). Animal experiments were approved by the University of Liverpool (UoL) ethics committee and performed under a licence granted by the UK home office under the Animals (Scientific Procedures) Act 1986. Reporting of experiments is in line with the ARRIVE guidelines. Animals were closely observed for any side effects and were culled at the end of experiments (day 14 post injection) without observing a noticeable tumor mass.

## Methodology

### Synthesis of super-paramagnetic iron oxide (maghemite) nanoparticles (SPIONs)

SPIONs were synthesized following the published protocol from Barrow *et al*.^19, 20^ Briefly, maghemite (SPIONs) were synthesized by mixing DEAE-dextran (0.05 g, Mw 40 kDa), ferric chloride hexahydrate (0.03 g), and ferrous chloride tetrahydrate (0.015 g) in 25 mL water with non-magnetic (polytetrafluoroethylene) stirring under an air-tight connection. The mixture was purged with nitrogen on ice (30 minutes) and ammonium hydroxide (1 mL, 28 - 30%) was added dropwise (in 2 min) under stirring (200 ± 5 rpm). This was followed by mixture transfer from ice to an oil bath (pre-set at 60 °C) and temperature elevation to 80 °C under nitrogen (in 15 min) and then left at 80 °C for 1 h. Afterwards, the product was brought under air, and refluxed for 5 hrs at 110 °C. SPIONs were purified at room temperature by dialysis (100 kDa membrane) until the pH became neutral (7). After concentrating by spin filter (1 - 2 mL), the sample was passed through size exclusion beads (dextran-based G-100 Sephadex^®^) to remove excess free polymer. Finally, the particles were washed thrice with deionized water by spin filteration. SPIONs were passed through 0.22 µm polyethersulfone (PES) membrane for sterilisation before use in experiments and saved at 4 °C.

### Synthesis of SPIONs encapsulating PEM capsules

SPIONs were encapsulated by a modified co-precipitation method,^28, 34^ and LbL assembly of oppositely charged layers of polymers and SPIONs around co-precipitated cores. PEM capsules consisting of 32 layers [(PSS/PAH/PSS/SPIONs)7/(PSS/PAH)2] of non-biodegradable polymers, i.e., PSS and PAH, and SPIONs were fabricated around sacrificial template cores comprising of calcium carbonate incorporating SPIONs and high molecular weight dextran. In order to co-precipitate SPIONs and dextran with calcium carbonate microparticles, solutions of calcium chloride and sodium carbonate were mixed under vigorous stirring in the presence of SPIONs and dextran at room temperature. For this in a glass vial, 4.2 mL mixture (0.3 mL SPIONs; 0.95 mg/mL, 1 mL dextran; 6.5 mg/mL, and 2.9 mL water) was added to 3 mL of calcium chloride solution (0.33 M). 3 mL of sodium carbonate solution (0.33 M) was quickly added to the above mixture and continuously stirred for 30 s, under vigorous magnetic stirring (1500 rpm) followed by keeping the reaction contents without agitation for 2 min at room temperature. The resulting calcium carbonate particles incorporating SPIONs and dextran were washed thrice with deionized water and used for LbL assembly of oppositely charged polyelectrolytes (2 mg/mL in 0.5 M NaCl) and SPIONs (0.5 mL SPIONs; 0.95 mg/mL, 4 mL water, and 0.5 mL; 0.5 M sodium chloride). The alternating layers of negatively and positively charged polymers, and SPIONs were electrostatically deposited around the charged sacrificial microparticle template cores following an established protocol.^48, 49^ For this the microparticles were alternatively immersed inside the solutions of either layer (5 mL; PSS, PAH, or SPIONs), exposed to sonication for short time (1 - 3 min) and kept under agitation/shaking for 13 min. After deposition of each layer the particles were washed thrice with deionized water to remove excess reactants (polymers/SPIONs, etc.). Finally, the cores of PEM capsules were dissolved by complexion of calcium ions with EDTA (5 mL, 0.2 M, pH 6.5; 30 min incubation), washed with deionized water, and stored at 4 °C after addition of 10.5 mL water for further use. The capsules diameter was ~ 2 µm as observed from TEM analysis. Further in the manuscript these PEM capsules incorporating SPIONs are called encapsulated SPIONs.

For SQUID measurements, free and encapsulated SPIONs were freeze-dried and SQUID measurements, TEM, pXRD (free SPIONs), zeta size and potential measurements were performed as mentioned in the characterization section.

### Ferrozine assay and ICP-OES based Fe quantification

The samples (particles and particle-labelled cells) were dissolved in 1.2 M HCl. Fe standard curves were made from the known concentrations of Fe and from these standard curves the unknown concentrations of Fe in the samples were determined.

For ferrozine assay, the Fe standard (50 µL, 4 µg in 1.2 M HCl) was serially diluted in HCl (1.2 M) followed by addition of 50 µL water so that final Fe concentration in standards was 2, 1, 0.5, 0.25, 0.125, 0.0625, 0.031 µg in HCl (0.6 M; 100 µL). Blank was 100 µL HCl (0.6 M). In the samples with unknown Fe content, 50 µL HCl (1.2 M) was added and samples, Fe standards and blank were placed on a heat block at 65 °C for 2 hrs. After heat-assisted acid dissolution, the samples and standards were cooled down to room temperature and centrifuged to collect the condensation products along the walls of tubes. HCl concentration was adjusted to 0.6 M by adding water. 30 µL ferrozine reagant was added to 100 µL of each sample including standards and blank, mixed and left at room temperature for 30 min. Their absorption was measured at 590 and 780 nm using a plate reader. For background correction the absorbance at 780 nm was subtracted from the absorbance at 590 nm and blank was subtracted from all values. A standard curve for Fe content was plotted and was used to quantify Fe contents in the samples with unknown Fe concentrations.

(Note: the ferrozine reagent was prepared by dissolving 2.4 g ammonium acetate, 2.2 g ascorbic acid, 0.02 g ferrozine, and 0.02 g neocuproine (dissolved in small volume of ethanol) in 6.25 mL water, mixed, frozen in small aliquotes and protected from light).

For ICP-OES Fe quantifications after dissolution with 1.2 M HCl, the samples were diluted in water so that final acid content will not exceed 5%. ICP-OES measurements were performed using 3 readings per wavelength and 11 wavelengths per sample. The concentration determination was performed using calibration curve for Fe consisting of 6 measurement points of freshly prepared Fe concentrations (0 - 10 µg/mL) derived from iron standard solutions from Inorganic Ventures (100 mg/L).

### Cell viability measurements

CellTiter-Glo^®^ reagent was used for cell viability measurements. For this assay, mMSCs were seeded in 96-well transparent bottomed plates at 10,000 cells per well in 100 µL complete cell growth medium (DMEM supplemmented with 10% FBS, 1% P/S, and 1% L-glutamine). After 24 hrs cells were incubated with free and encapsulated SPIONs suspended in fresh cell culture medium for 24 hrs. Serial dilution of particles was performed in complete cell growth medium. Each condition was applied in triplicate. After incubation with particles, the cells were washed with PBS and 100 µL fresh cell growth medium was added to each well of assay plates, followed by the addition of 20 µL CellTiter-Glo^®^ reagent. The plates were shaken at 600 - 700 rpm in an orbital shaker for 1 min (to complete cell lysis), incubated at room temperature for 5 min and 100 µL mixture contents from each well of the plates were transfered in opaque bottomed 96 well plates. The luminescence from each well of 96 well plate was measured by a Fluostar Omega (BMG Labtech) plate reader. Viability was expressed as % of the untreated control.

### Phantoms for MR measurements

Phantoms for MR measurments were prepared in 1% low melting agarose.

### 1- Phantoms of particles

Low melting agarose was dissolved in water at 65 °C to get a clear 2% solution and placed at 40 °C during mixing and addition of particles. The solutions of SPIONs (free and encapsulated) were diluted in water (serial dilution) to get 0.5, 0.25, 0.125, 0.0625, and 0.03125 mM Fe concentrations. These diluted solutions of particles were mixed with the gel at 40 °C (1:1 dilution by volume; 100 µL of each sample was mixed throughly with 100 µL gel without bubble formation in 200 µL polypropylene tubes) to get final Fe content 0.25, 0.125, 0.0625, 0.03125, and 0.0156 mM in 1% agarose. 2% agarose was mixed with water (1:1 dilution) to serve as control. These samples were mounted in 1% agarose containing holders, stored at 4 °C for 24 hrs and analyzed by MRI. The MR measurement parameters are listed in Table SI.1. The solution relaxivity of the particles entrapped in phantoms was calculated from their relaxation times which were acquired by multi gradient echo sequences (MGES) and the imaging was performed with a fast low angle shot (FLASH) sequence.

### 2- Phantoms of labelled cells

In 6 well plates mMSCs were seeded (2 × 10^5^ cells per well) in 2 mL complete growth media. After 24 hrs the free and encapsulated SPIONs were added at 15, 20, and 25 capsules per cell (equivalent to a dose of 13.1, 17.5, and 21.8 pg Fe in free SPIONs per cell) considering the seeded number of cells. After 24 hrs of incubation, cells were washed twice with PBS and trypsinized. Particle-labelled cells were centrifuged at 500 rcf for 5 min and the supernatant was removed. The cell pellets were resuspended in 4% PFA, mixed without bubble formation and fixed cells were transfered in small polyproplylene tubes (100 µL for each condition) centrifuged and maxiumum supernatant was removed. The cell pellets were resuspended in PBS and mixed with 2% low melting agarose (1:1 dilution). 3 × 10^5^ fixed cells were poured on the top of already solidifed 1% agarose gel. The samples were mounted in 1% agarose containing holders and imaged with a fast low angle shot (FLASH) sequence. 19 images per sample were captured. Their T2 and T2* (T2-Star) relaxation times were measured with MGES. Imaging parameters are listed in Table SI.2.

For ferrozine based Fe quantification, the particles labelled cells (after washing and trypsinization) were counted, centrifuged and dissolved in HCl (1.2 M) after removing maximum supernatant.

### Luc^+^ activity of mMSCs

The cells were monitored for Luc^+^ activity before *in vivo* experiments. For this mMSCs were seeded in 96 well plates in triplicate with serial dilutions (10,000, 5000, 2500, 1250, 625, 312, 156, 78, 39, 20, 7, 3, 2, 1, and 0 cells per well) in 100 µL complete growth media and cultured at constant (5%) supply of CO^2^ for 24 hrs at 37 °C. After refreshing the growth media, the plates were placed in IVIS spectrum imaging system, and noted the baseline bio-luminescnce signals. Latter, their growth media was replaced by fresh growth media (100 µL) containing D-luciferin (15 µg/mL), left at room temperature for 15 min and their bio-luminescence was recorded and expressed as radiance (photons/second/cm^2^/steradian [p/s/cm^2^/sr]).

### Cell labelling and administration in mice

mMSCs transfected with lentiviral vector (Luc^+^) were IC injected in mice. The vector details are described in a recent study.^12^ To label cells with particles, 1.5 × 10^6^ mMSCs were seeded in 100 mm tissue culture dishes and incubated for 24 hrs at 37°C in humidified atmosphere with 5% CO^2^. Free (33.72 pg Fe/cell) and encapsulated (@15 capsules per cell equivalent to a dose of 13.1 pg Fe/cell) SPIONs were added to the cells considering the seeded number of cells and incubated for 24 hrs. The cells were washed twice with PBS, trypsinized, kept on ice (after removing trypsin), and resuspended in ice cold PBS after cell counting. To observe the morphology of cells after particles labelling, some of the trypsinized cells were re-seeded and imaged after 3 days. 10^6^ cells in 100 µL PBS (having ~5 and ~4 pg Fe/cell for free and encapsulated SPIONs, respectively) were IC injected into the left ventricles of female BALB/c mice under ultrasound guided injection. Details of IC injections is provided in a recent study.^11^ Briefly, mice were anesthetized with a mixture of isoflurane and oxygen, subcutaneous (SC) injected with an analgesic buprenorphine (0.1 mg/kg body weight) and were positioned supine above a heated platform. Their fur was removed and the limbs and abdomen were taped after extension of body (to hold body in extended position and the skin over the chest taut). Ultrasound gel was applied to the chest, and ultrasound tranducer was positioned over the mice to have chest in view. Visualized the heart and labelled cells were injected inside the left cardiac ventricle with the help of 29G ½ inch insulin syringe. The cells were completely re-suspended, and were injected in a slow and well controlled manner. Soon after IC injections mice were imaged by MR under the same anaesthesia session and latter in groups of 4 by IVIS based BLI, on 0 (injections day), 1, 2, 6, 10, and 14 days post injections. Where possible mice were imaged by both imaging modalities under same anesthetic session. The mice were recovered from anaesthesia in a heat box set at 32 °C, and closely monitored for the signs of any adverse effects during the course of studies.

### *In vivo* MR imaging

The biodistribution of cells in the abdomenal region, i.e., kidneys, liver, spinal muscles, and spleen was imaged with a 9.4T horzontal bore MRI scanner (Bruker Avance III spectrometer) using a 40 mm transmit/receive volume coil. FLASH T2* weighted sequences were recorded and T2* maps were generated using MGES. A prescan before injections served as control to set baseline value for MR analysis. Regions of interests (ROIs) were drawn around the kidney cortices, medullae, livers, spleen and spinal muscles and T2* relaxation times were calculated from the T2* map images. MR acquisition parameters are listed in Table SI.3.

### Bioluminescence imaging (BLI)

BLI complementary to each MR scan was performed until the last imaging day (day 14 post injection). Mice received D-luciferin (150 mg/kg body weight) intraperitoneally and were imaged 15 min post injections by IVIS spectrum imaging system. Imaging was performed by 1 - 3 min luminescnce exposure and expressed as radiance (photons/second/cm^2^/steradian [p/s/cm^2^/sr]).

### ICP-OES based Fe quantification in mice organs

For quantification of Fe content in the brain, kidneys, liver, lungs, spleen, and heart, at the end of *in vivo* experiments mice were killed under terminal anesthesia followed by cervical dislocation. After dissection, the above mentioned organs were collected, washed with PBS and preserved in 70% ethanol. Organs collected from 3 untreated mice served as control. The organs in 70% ethanol were dipped in 96% ethanol, crushed with pestle and mortar and left overnight for freeze-drying. The dried crushed organs mass was weighed. 0.5 mL HCl (1.2 M) was added to the dried and crushed organs, kept inside an oven at 70 °C for 3 hrs and brought the reaction contents at room temperature. Water was added to each sample vial to get 10 mL final voulme. Samples were filtered through 0.2 µm filters to remove organ debris. Fe content in each sample was determined by ICP-OES. The dried weight of crushed organs noted before acid digestion was used for normalizing Fe content per dried weight of each organ.

### Conclusions

We have demonstrated that SPIONs can be encapsulated inside polymeric capsules while retaining their individual particle identities, and showed similar magnetization behavior as free particles. Although the solution relaxivity of free particles was very different from encapsulated ones, once internalized inside lysosomal compartments after cell uptake, this difference became negligible for similar concentrations (Fe) of particles. The encapsulation of particles has some potential advantages over free particles; e.g., (1) in case of cell labelling with small amount of particles, a higher uptake can be achieved, and (2) the addition of different types of particles in a single entity for multiplexed measurements is possible. Comparison of both formulations of SPIONs did not reveal any difference in the *in vivo* fate of particles upon cell death and was similar to what has been reported for particles directly injected in the blood stream of mice.

## Acknowledgements

We thank Dr Antonius Plagge for providing Luc^+^ mMSCs. The Electron Microscopy Unit, Biomedical Services Unit, and the Centre for Preclinical Imaging at the University of Liverpool are acknowledged for help and support. We are grateful for the support from BBSRC, EPSRC, and MRC-funded UK Regenerative Medicine Platform “Safety and Efficacy, focusing on Imaging Technologies Hub” (MR/K026739/1). SA acknowledges a Fellowship from the Marie Skłodowska-Curie (MultimodalCellTrack/705600). Pranab Mandal and Stephen Moss are acknowledged for SQUID and ICP-OES measurements, respectively.

